# Photoreceptor-derived FGF2 mediates a protective stress response without driving pathological retinal neovascularization in ischemic retinopathy

**DOI:** 10.64898/2026.08.19.745869

**Authors:** Zhifei Wu, Lulu Peng, Jiechun Wu, Huirong Xu, Yaling Liu, Dingqiao Wang, Lan Wang, Xingyu Huang, Guoming Zhang, Peng Wang, Wei Du

**Affiliations:** Department of Ophthalmology, The Eighth Affiliated Hospital, Sun Yat-Sen University, Shenzhen, China; Biological Laboratory of Hetao Cooperation Zone, The Eighth Affiliated Hospital, Sun Yat-sen University, Shenzhen, China; Department of Orthopedics, The Eighth Affiliated Hospital, Sun Yat-Sen University, Shenzhen, China; Shenzhen Eye Hospital, Shenzhen Eye Medical Center, Southern Medical University, Shenzhen, China

## Abstract

Fibroblast growth factor 2 (FGF2) is frequently induced during ischemic retinal injury and has traditionally been considered a pro-angiogenic factor based largely on studies using exogenous FGF2 administration. However, its endogenous cellular origin and physiological role remain incompletely understood. Here, we used single-cell transcriptomic analysis combined with spatial validation and rod photoreceptor-specific genetic approaches to define the endogenous role of FGF2 during oxygen-induced retinopathy (OIR). We identify rod photoreceptors as a major cellular source of ischemia-induced FGF2. Notably, *Fgf2* expression remained elevated during the regression of pathological neovascularization, revealing a temporal dissociation between neuronal stress responses and vascular remodeling. Single-cell analysis further showed that *Fgf2* induction occurred within a coordinated photoreceptor stress-response program involving endothelin 2 (*Edn2*) and B-cell lymphoma 3 (*Bcl3*). This transcriptional signature was independently reproduced in the N-methyl-N-nitrosourea (MNU)-induced photoreceptor degeneration model. Rod-specific deletion of *Fgf2* markedly increased photoreceptor apoptosis, indicating that endogenous FGF2 contributes to photoreceptor survival under ischemic stress. In contrast, neither genetic depletion nor overexpression of FGF2 altered pathological neovascularization or vaso-obliteration. Bidirectional manipulation of FGF2 further modulated the expression of representative stress-associated genes *Edn2*and *Bcl3,* supporting FGF2 involvement in this injury-response program. Finally, receptor expression analysis revealed relatively limited endothelial expression of *Fgfr1-Fgfr4* compared with VEGF receptors, suggesting a cellular basis for the distinct effects of endogenous FGF2 and VEGF signaling. Together, these findings identify endogenous retinal FGF2 as a photoreceptor-derived survival factor that is induced during stress but is insufficient to drive pathological angiogenesis. These results support a model in which neuronal adaptation and vascular remodeling represent partially distinct responses during ischemic retinal injury.

## Introduction

Ischemic retinopathies, including retinopathy of prematurity (ROP), proliferative diabetic retinopathy (PDR), and retinal vein occlusion (RVO), remain leading causes of irreversible vision loss worldwide ^[1]^ . A central pathological event in these disorders is retinal ischemia, which drives hypoxia-induced pathological neovascularization ^[2]^. Intravitreal anti-vascular endothelial growth factor (VEGF) therapy has substantially improved clinical outcomes by suppressing pathological vessel growth; however, anatomical vascular improvement does not always translate into complete visual recovery ^[3–6]^. This clinical dissociation suggests that neuronal injury and pathological vascular remodeling may be governed by partially distinct biological programs, highlighting the importance of understanding endogenous mechanisms that preserve neuronal integrity during ischemic stress.

FGF2 is a pleiotropic growth factor involved in diverse biological processes, including development, tissue repair, stem cell maintenance, neuronal survival, and angiogenesis^[7,8]^. In multiple experimental systems, exogenous FGF2 potently stimulates endothelial proliferation and neovascularization, establishing it as a prototypical pro-angiogenic molecule^[9,10]^. Consistent with this view, elevated FGF2 levels have been detected in vitreous samples from patients with proliferative diabetic retinopathy and in experimental models of retinal ischemia^[11,12]^.Nevertheless, accumulating evidence also indicates that FGF2 possesses neuroprotective properties, with early studies demonstrating that exogenous FGF2 delays photoreceptor degeneration and promotes retinal neuronal survival in models of inherited retinal degeneration and light-induced retinal injury ^[13–15]^ . Despite extensive investigation, however, it remains unresolved whether endogenous retinal FGF2 functions as a physiologically relevant regulator of pathological neovascularization, particularly given the fundamental differences between exogenous administration and endogenous ligand activity within the native tissue microenvironment.

Previous genetic studies have provided important evidence against a major role for endogenous FGF2 in pathological retinal neovascularization. Constitutive deletion of *Fgf2*has little effect on pathological neovascularization in the mouse OIR model, whereas photoreceptor-targeted FGF2 overexpression fails to induce spontaneous retinal angiogenesis ^[16]^. These findings argue against a dominant angiogenic role for endogenous FGF2 under ischemic conditions. Nevertheless, both constitutive knockout and conventional transgenic overexpression strategies are subject to interpretational limitations, including potential developmental compensation, systemic effects, and non-physiological expression levels^[17]^, leaving the cell-type-specific contribution of retinal FGF2 unresolved. Furthermore, recent studies suggest that FGF2 produced by activated microglia may indirectly influence vascular responses through inflammatory pathways ^[18,19]^, highlighting the complexity of endogenous FGF2 signaling. Thus, defining the cellular source and functional context of endogenous FGF2 remains essential for resolving its physiological role during retinal ischemic injury.

Unlike many secreted growth factors, FGF2 lacks a canonical signal peptide and is released through unconventional secretion mechanisms, making protein localization alone insufficient to reliably identify its endogenous cellular source ^[20]^. Recent advances in single-cell transcriptomics offer a powerful approach to resolve ligand expression at cellular resolution ^[21]^. Importantly, the biological effects of secreted factors depend not only on their cellular origin but also on whether potential target cells express the appropriate receptors. Therefore, defining both the cellular source of FGF2 and the receptor landscape of potential target cells is critical for understanding its physiological function in the ischemic retina.

Beyond vascular remodeling, accumulating evidence indicates that photoreceptors activate intrinsic stress-response programs in response to retinal injury and degeneration. Transcriptomic analyses across multiple models of photoreceptor degeneration have revealed shared induction of injury-associated genes, including *Edn2, Bcl3,* and *Fgf2,* suggesting the activation of conserved molecular responses during photoreceptor stress^[22–25]^. Among these factors, *Edn2*functions as a photoreceptor-derived stress signal that mediates communication with surrounding retinal cells and contributes to photoreceptor adaptation and survival in inherited degeneration models ^[26,27]^. *Bcl3,* a transcriptional regulator that modulates NF-κB-dependent gene expression, has also been implicated in neuronal protection and retinal injury responses ^[27,28]^, although its role in photoreceptor stress adaptation remains unclear. A central question is whether stressed photoreceptor-derived FGF2 represents a functional component of an intrinsic adaptive response or merely an injury-associated bystander. Furthermore, whether FGF2 induction parallels vascular remodeling or reflects a sustained neuronal stress response during ischemic retinal injury remains to be systematically addressed.

Here, we sought to define the cellular origin and endogenous function of FGF2 during ischemic retinal injury by integrating single-cell transcriptomics, spatial validation, temporal profiling, and cell-type-specific genetic manipulation. We identify rod photoreceptor-derived FGF2 as a component of an injury-response program that persists beyond pathological neovascular regression and supports photoreceptor survival while remaining dispensable for pathological neovascularization. These findings reveal how cellular origin and receptor availability jointly shape growth factor function in the ischemic retina.

## Results

### Rod photoreceptors are the major source of sustained FGF2 induction during ischemic retinal injury

To define the cellular source and temporal dynamics of endogenous FGF2 during ischemic retinal injury, we first reanalyzed single-cell RNA-sequencing datasets from normoxic and OIR retinas ^[29]^.*Fgf2*transcripts were detected predominantly in rod photoreceptors, whereas substantially lower expression was observed across other major retinal cell populations, including Müller glia, cone photoreceptors, microglia and astrocytes (Fig. 1A, B). These data identified rod photoreceptors as a major cellular source of ischemia-induced *Fgf2* expression.

**Figure 1.**
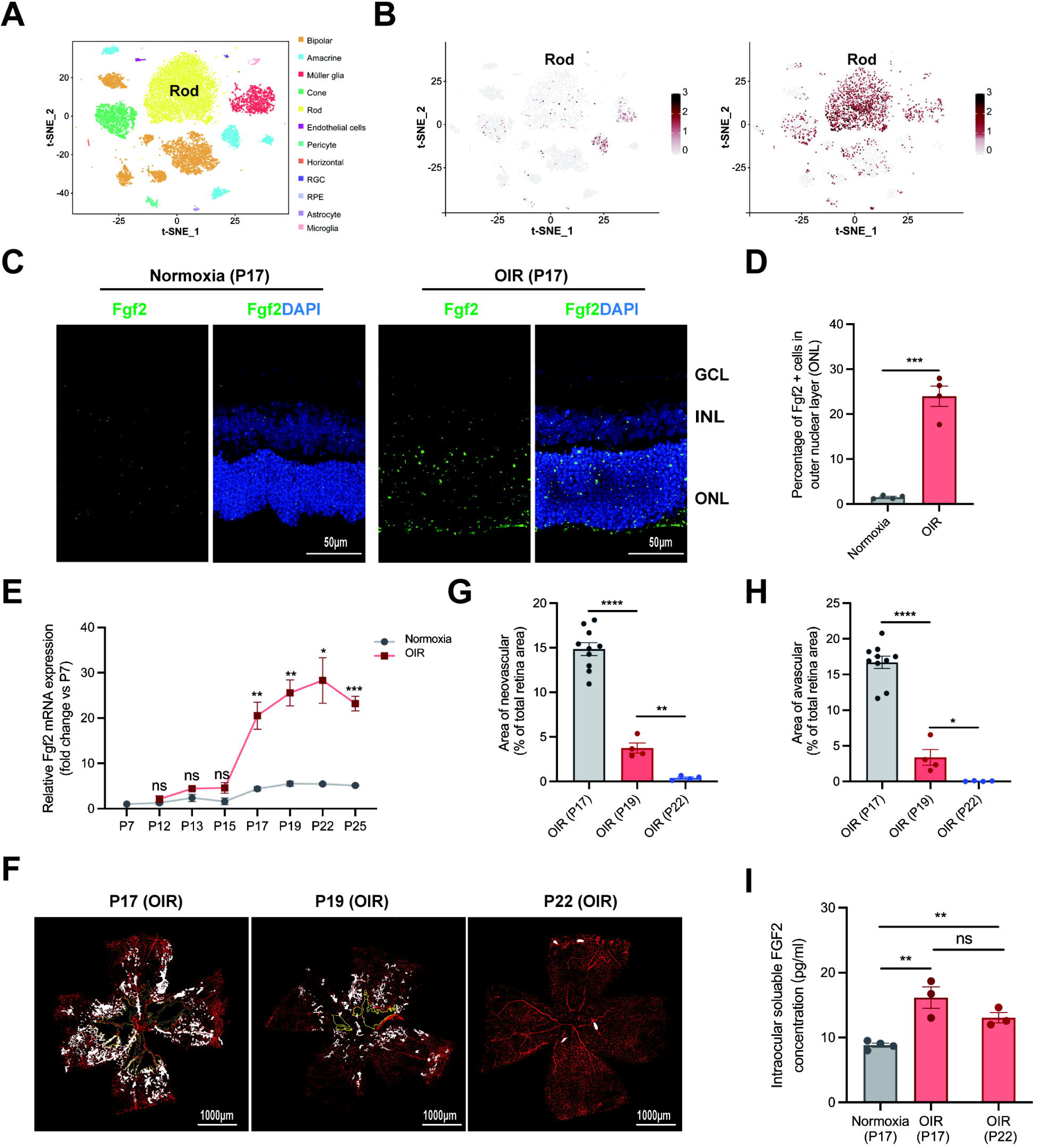
FGF2 is predominantly induced in rod photoreceptors and remains elevated during the regression of pathological neovascularization. (A) t-SNE visualization of single-cell RNA sequencing data showing the major retinal cell populations identified in the dataset. (B) t-SNE feature plots showing *Fgf2* expression in normoxic and OIR retinas at P17. (C) Representative RNAscope images showing *Fgf2* expression in normoxic and OIR retinas at P17. *Fgf2* transcripts were detected by RNAscope and visualized as *Fgf2* signal alone and in combination with DAPI counterstaining. Scale bar, 50 μm. (D) Quantification of the percentage of *Fgf2-* positive cells among ONL cells in normoxic and OIR retinas at P17. Data are presented as mean ± SEM; *n* = 4 mice per group. \*\*\**P*<0.001, two-tailed unpaired Student’s *t*-test. (E) Temporal expression of retinal *Fgf2* mRNA in normoxic and OIR mice from P7 to P25, determined by quantitative RT-PCR. Data are presented as mean ± SEM; *n*= 3 mice per time point. Statistical significance between OIR and normoxic groups at each individual time point was determined by unpaired two-tailed Student’s *t*-test, followed by Benjamini-Hochberg false discovery rate (FDR) correction for multiple comparisons across all time points. ns, not significant; \**P*<0.05, \*\**P*<0.01, \*\*\**P*<0.001. (F) Representative IB4-stained retinal flat mounts from OIR mice at P17, P19, and P22. White-filled areas indicate pathological neovascular tufts, and yellow outlines indicate avascular areas. Scale bar, 1000 μm. (G, H) Quantification of pathological neovascularization (G) and avascular (H) areas in OIR retinas at P17, P19, and P22. Data are mean ± SEM; *n*≥ 4 mice per time point. Statistical significance was determined by one-way ANOVA with Tukey’s multiple-comparison test. ns, not significant; \**P*< 0.05, \*\**P*< 0.01, \*\*\*\**P*< 0.0001. (I) Soluble FGF2 protein levels in ocular fluids (aqueous and vitreous humor) collected from normoxic P17, OIR P17, and OIR P22 mice, determined by ELISA. Data are presented as mean ± SEM; *n*≥ 3 pooled samples per group (each pool from 6-8 mice). Statistical significance was determined by one-way ANOVA with Tukey’s multiple-comparison test. ns, not significant; \*\**P*< 0.01.

To corroborate the cellular distribution of Fgf2 transcripts, we performed RNAscope in situ hybridization. *Fgf2-*positive cells were rarely detected in the outer nuclear layer (ONL) of normoxic retinas, whereas OIR retinas exhibited a marked increase in *Fgf2-*positive cells within the ONL (Fig. 1C, D). Quantification confirmed a significant increase in the proportion of *Fgf2-*positive ONL cells following OIR. These results, together with the single-cell transcriptomic data, establish rod photoreceptors as the predominant cellular source of ischemia-associated FGF2 expression.

Having established the cellular origin of *Fgf2* expression, we next assessed its temporal relationship with pathological retinal vascular remodeling. As expected, pathological neovascularization was prominent at P17 and progressively declined at later OIR time points (Fig. 1F,G). In contrast, retinal *Fgf2* expression began to increase after P15 and remained elevated during the period in which pathological neovascularization was undergoing regression (Fig. 1E). Consistent with this pattern, intraocular FGF2 protein levels were increased in the ocular fluid of OIR mice at both P17 and P22 compared with normoxic controls, with no significant difference between OIR P17 and OIR P22 (Fig. 1I). Thus, retinal FGF2 expression was not temporally restricted to the phase of maximal pathological neovascularization but remained elevated during the subsequent vascular regression phase.

These findings indicate that ischemic retinal injury induces sustained FGF2 expression predominantly in rod photoreceptors and that the temporal profile of FGF2 induction is not tightly coupled to the progression of pathological neovascularization.

### FGF2 induction is associated with a coordinated photoreceptor injury-response program

The persistence of FGF2 expression during the regression phase of pathological neovascularization led us to investigate whether *Fgf2* induction was associated with a broader photoreceptor stress-response program. Analysis of *Fgf2* expression together with a panel of previously reported photoreceptor injury-response genes showed a shared induction pattern involving *Edn2, Bcl3, Fos,* Jun family members, *Atf3,* and *Mt1* in OIR photoreceptors (Fig. 2A).

**Figure 2.**
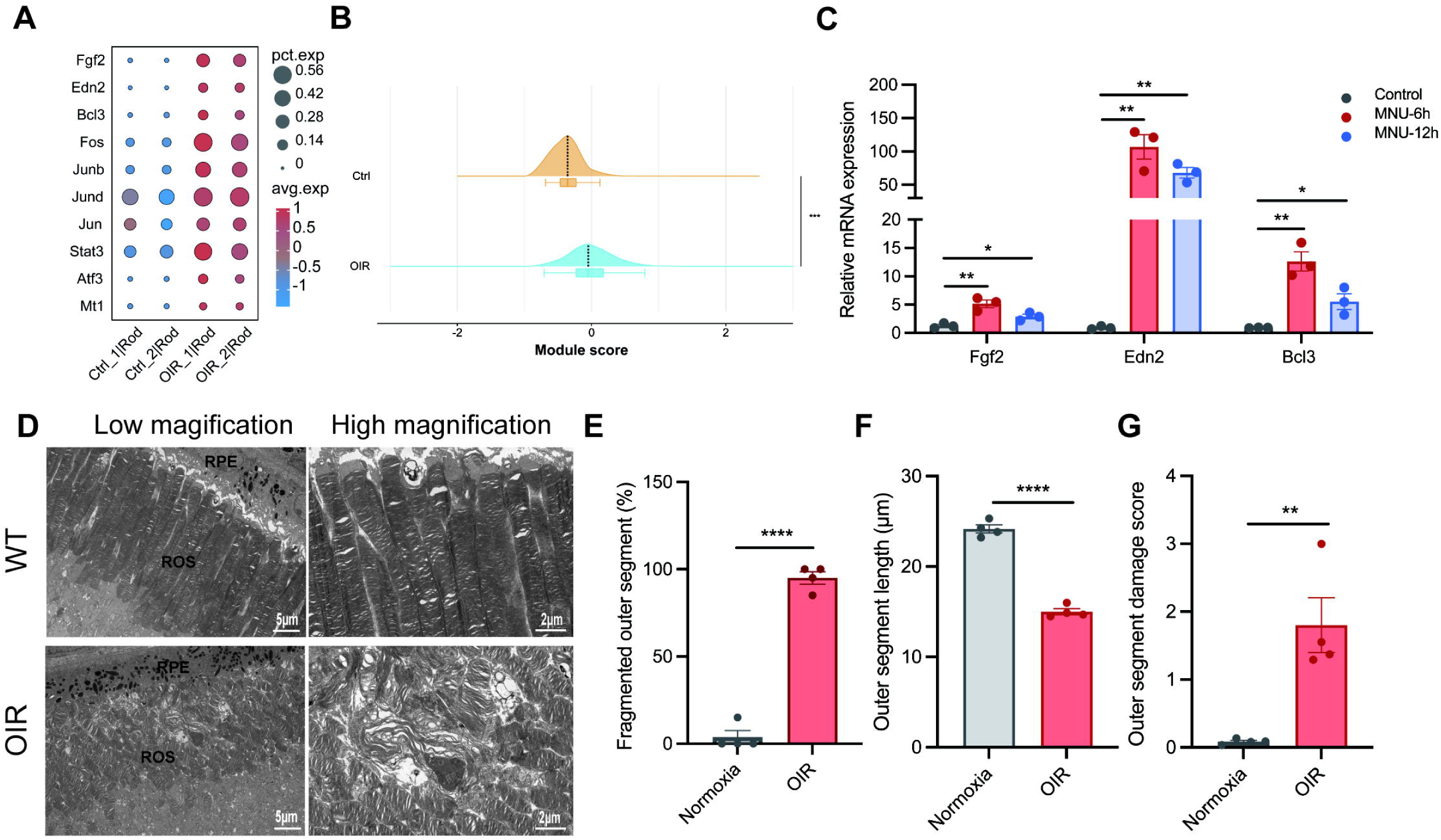
FGF2 is induced as part of a conserved photoreceptor injury-response program. (A) Dot plot showing expression of injury-response genes (*Fgf2*, *Edn2*, *Bcl3*, *Fos*, *Junb*, *Jund*, *Jun*, *Stat3*, Atf3, and *Mt1*) in rod photoreceptors from two independent normoxic and two independent OIR single-cell RNA sequencing datasets. Dot size represents the percentage of expressing cells and color indicates average normalized expression. (B) Ridge plot showing the distribution of the module score calculated from the predefined injury-response gene set in control and OIR rod photoreceptors. \*\*\**P*< 0.001. (C) Quantitative RT-PCR analysis of retinal *Fgf2*, *Edn2*, and *Bcl3* mRNA levels in MNU-treated mice at 0, 6, and 12 h post-injection. Data are presented as mean ± SEM; *n*= 3 mice per time point. Statistical significance was determined by one-way ANOVA with Dunnett’s multiple-comparison test. \**P*< 0.05, \*\**P*< 0.01. (D) Representative transmission electron microscopy images of photoreceptor outer segments in normoxic and OIR retinas at P17. Left panels show low-magnification overviews of the outer retinal layers; right panels show high-magnification views of the outer segment ultrastructure.ROS, rod outer segment; RPE, retinal pigment epithelium. Scale bars: 5 μm (low magnification) and 2 μm (high magnification). (E-G) Quantification of outer segment fragmentation (E), outer segment length (F), and ultrastructural damage score (G), respectively. *n*= 4 mice per group. Data are presented as mean ± SEM. \*\**P*<0.01, \*\*\*\**P*<0.0001, two-tailed unpaired Student’s t-test.

To assess this response at the level of a broader transcriptional pattern, we generated a composite injury-response module score based on the predefined injury-response gene set. Rod photoreceptors from OIR retinas exhibited a significant increase in the module score compared with control photoreceptors (Fig. 2B). Thus, *Fgf2* induction occurred within a broader photoreceptor injury-response state rather than as an isolated transcriptional event.

Whether this injury-response signature was restricted to ischemic injury remained unclear. To test this possibility, an independent model of chemically induced photoreceptor degeneration using MNU recapitulated the induction pattern of these genes ^[30]^. Following MNU administration, retinal expression of *Fgf2, Edn2,* and *Bcl3* increased during the early phase of photoreceptor injury (Fig. 2C). The shared induction of these genes in a mechanistically distinct model of photoreceptor degeneration indicated that this response was not unique to the vascular pathology of OIR.

Consistent with the transcriptional evidence of photoreceptor injury, transmission electron microscopy demonstrated substantial structural abnormalities in photoreceptor outer segments during OIR. Compared with normoxic controls, OIR retinas exhibited outer segment fragmentation, reduced outer segment length, and increased ultrastructural damage (Fig. 2D-G, Fig. S3).

Collectively, these observations demonstrate that FGF2 induction occurs within a broader photoreceptor injury-response state that is also observed in an independent model of photoreceptor degeneration.

### Photoreceptor-derived FGF2 supports photoreceptor survival without altering pathological retinal angiogenesis

The association of *Fgf2* with photoreceptor injury-response genes raised the question of whether endogenous photoreceptor-derived FGF2 influences the pathological and neuronal consequences of ischemic retinal injury. To address this question functionally, we performed complementary loss- and gain-of-function analyses. Rod photoreceptor-specific deletion of *Fgf2*was achieved by crossing *Fgf2*^fl/fl^ mice with Rho-Cre mice. Efficient reduction of retinal *Fgf2* expression following rod-specific deletion was confirmed by quantitative RT-PCR (Fig. 3A, B). For gain-of-function studies, we used AAV8-Rho-*Fgf2* to increase FGF2 expression in rod photoreceptors. The efficiency and specificity of the rod-targeted approach were validated molecularly (Fig. 3A, C; Fig. S1). To complement rod-specific overexpression and test for broader effects, we additionally employed AAV2-CMV-*Fgf2* to achieve pan-retinal FGF2 overexpression (Fig. S2A).

**Figure 3.**
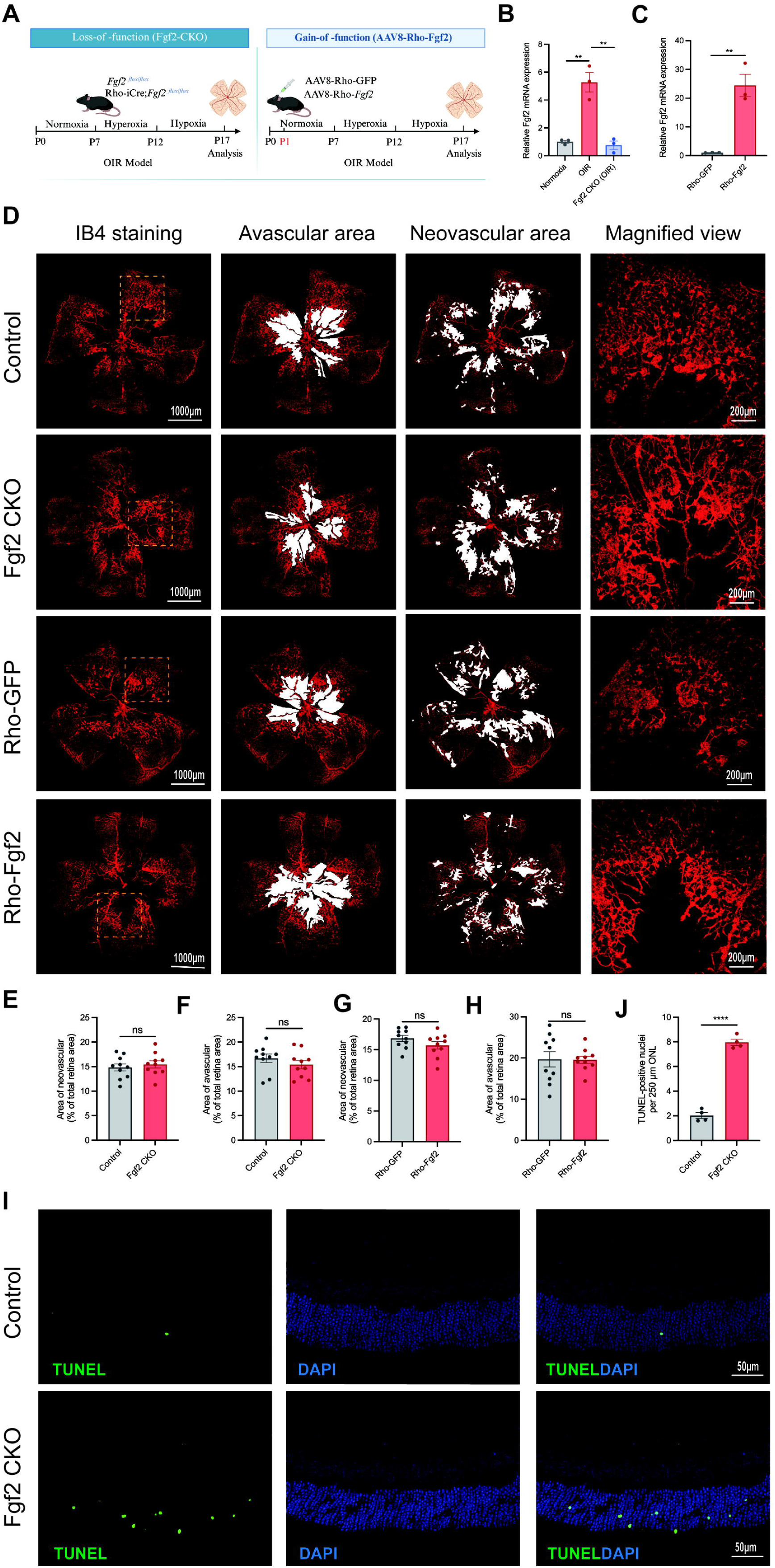
Endogenous FGF2 supports photoreceptor survival but does not alter pathological retinal angiogenesis. (A) Schematic illustration of the strategies used for rod photoreceptor-specific *Fgf2* conditional knockout and *Fgf2* overexpression. (B) Validation of rod photoreceptor-specific *Fgf2* deletion in OIR retinas at P17 by quantitative RT-PCR. Normoxic P17, OIR P17, and OIR P17 with conditional knockout of *Fgf2* are shown. Data are presented as mean ± SEM; *n*= 3 mice per group. Statistical significance was determined by one-way ANOVA with Tukey’s multiple-comparison test. \*\**P*< 0.01. (C) Validation of rod photoreceptor-specific *Fgf2* overexpression by quantitative RT-PCR following subretinal injection of AAV8-Rho-*Fgf2.* Data are presented as mean ± SEM; *n*= 3 mice per group. \*\**p*< 0.01, two-tailed unpaired Student’s *t*-test. (D) Representative IB4-stained retinal flat mounts from control, rod-specific *Fgf2* CKO, Rho-GFP, and Rho-*Fgf2* overexpression mice at P17, together with corresponding magnified views. Scale bars: 1000 μm (low magnification) and 200 μm (high magnification). (E-H) Quantification of retinal pathological neovascularization (E, G) and avascular areas (F, H) in OIR mice at P17 following rod-specific *Fgf2* CKO or *Fgf2* overexpression. Data are presented as mean ± SEM; *n*= 10 mice per group. Statistical significance was determined by unpaired two-tailed Student’s *t*-test. ns, not significant. (I) Representative TUNEL staining of retinal sections from control and *Fgf2* CKO mice at P17. Scale bar: 50 μm. (J) Quantification of TUNEL-positive nuclei per hot-spot field in the ONL. Hot spots were defined as the two most densely labeled microscopic fields per section, selected after surveying the entire ONL. At least three non-adjacent sections per mouse were analyzed, and the average count per hot-spot field was calculated as one biological replicate. Data are presented as mean ± SEM; *n*= 4 mice per group. Statistical significance was determined by unpaired two-tailed Student’s *t*-test. \*\*\*\**P*< 0.0001. Quantifications were performed in a blinded manner.

Functional assessment indicated that manipulation of photoreceptor-derived FGF2 did not significantly affect pathological vascular remodeling (Fig. 3D, E, G). Similarly, neither manipulation significantly changed the extent of avascular area (Fig. 3F, H). Consistently, pan-retinal FGF2 overexpression driven by the CMV promoter also failed to alter pathological neovascularization or vaso-obliteration (Fig. S2C-E). These results indicate that bidirectional manipulation of FGF2, achieved through rod-specific or ubiquitous retinal expression, did not produce a significant effect on the major vascular phenotypes of the OIR model.

In contrast, analysis of neuronal outcomes revealed a distinct phenotype. Rod-specific *Fgf2* deletion resulted in a marked increase in TUNEL-positive nuclei within the ONL of OIR retinas (Fig. 3I, J). Thus, loss of endogenous photoreceptor-derived FGF2 increased photoreceptor apoptosis during ischemic retinal injury. A functional dissociation emerges between the neuronal and vascular effects of photoreceptor-derived FGF2: under the conditions examined, endogenous FGF2 supports photoreceptor survival without appreciably altering pathological neovascularization or vaso-obliteration.

### FGF2 is functionally associated with representative components of the photoreceptor injury-response program

Given the association between *Fgf2* and the photoreceptor injury-response signature, we sought to determine whether FGF2 was functionally linked to representative components of this program. Because *Edn2* and *Bcl3* have established roles in photoreceptor stress adaptation and survival, we selected these genes as representative molecular readouts of the broader injury-response state.

At the single-cell level, *Fgf2* expression was positively correlated with *Edn2* and *Bcl3* expression in rod photoreceptors (Fig. 4A, B). These associations suggest that *Fgf2* expression occurs within the same transcriptional context as these stress-associated genes during OIR. Bidirectional manipulation of FGF2 levels demonstrated corresponding changes in *Edn2*and *Bcl3* expression. Rod-targeted FGF2 overexpression increased retinal *Edn2*and *Bcl3* mRNA levels compared with control AAV treatment (Fig. 4C). Conversely, rod-specific *Fgf2* deletion reduced the induction of *Edn2* and *Bcl3* in OIR retinas (Fig. 4D). Thus, bidirectional manipulation of FGF2 was associated with corresponding changes in the expression of these representative components of the photoreceptor injury-response program.

**Figure 4.**
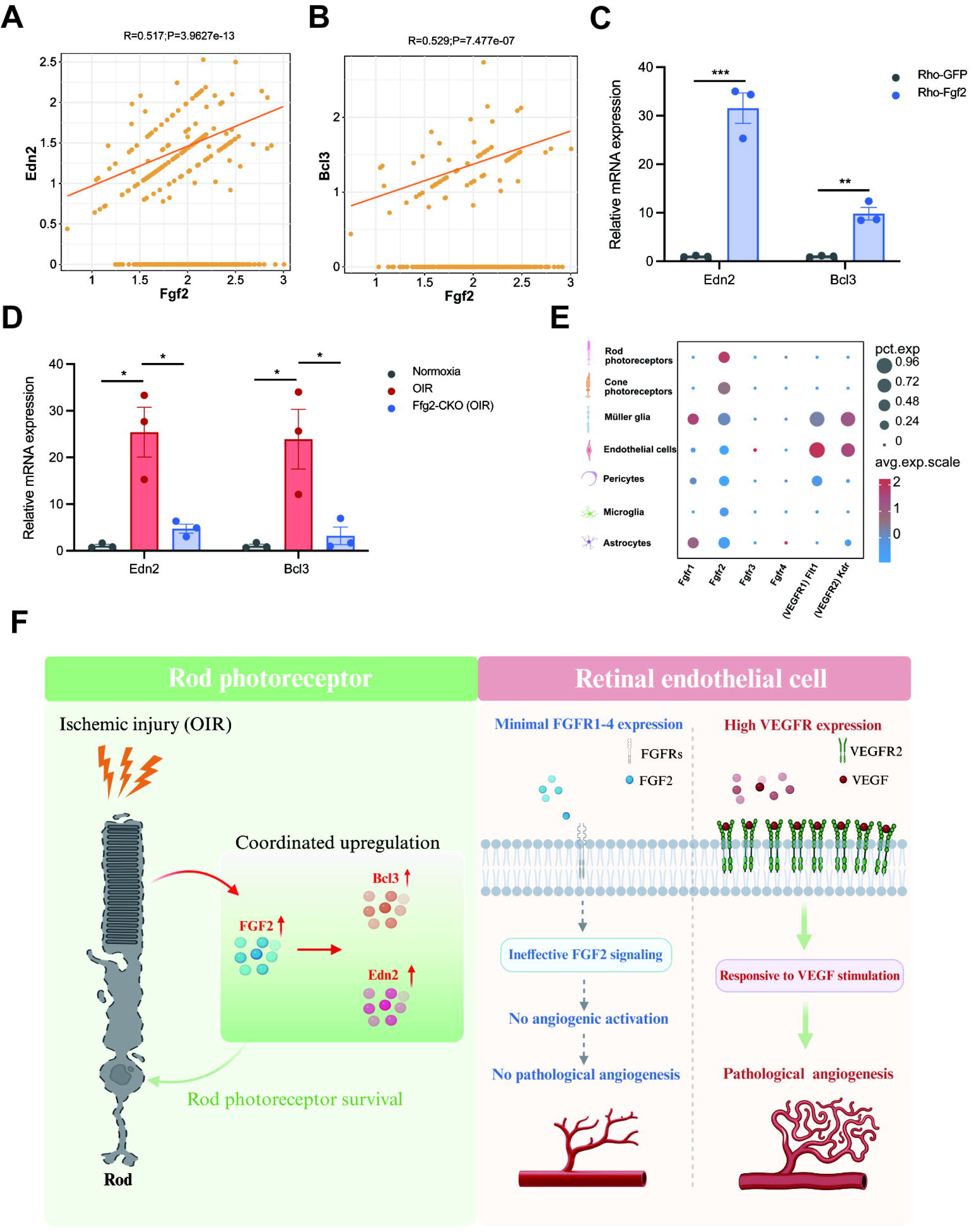
FGF2 is associated with a photoreceptor injury-response program but is uncoupled from endothelial angiogenic signaling. (A, B) Pearson correlation analysis of *Fgf2* with *Edn2*(A) and *Bcl3* (B) expression in rod photoreceptors (scRNA-seq). Double-negative cells were excluded. *Fgf2* positively correlated with *Edn2*(R = 0.517, *P*= 3.97 × 10^-13^) and *Bcl3* (R = 0.529, *P*= 7.48 × 10^-7^). (C) Quantitative RT-PCR analysis of *Edn2*and *Bcl3* mRNA expression in retinas of control (Rho-GFP) and *Fgf2* overexpression mice. Data are mean ± SEM; n = 3 mice per group. Statistical significance was determined by unpaired two-tailed Student’s t-test. \*\**P*<0.01, \*\*\**P*<0.001. (D) Quantitative RT-PCR analysis of *Edn2* and *Bcl3* mRNA expression in retinas of normoxic P17, OIR P17, and *Fgf2* CKO-OIR P17 mice. Data are mean ± SEM; n = 3 mice per group. Statistical significance was determined by one-way ANOVA with Dunnett’s multiple-comparison test. \**P*<0.05. (E) Dot plot showing expression of *Fgfr1-Fgfr4* together with canonical VEGF receptors (*Flt1* and *Kdr*) across major retinal cell populations. Retinal endothelial cells exhibited minimal *Fgfr* expression but abundant Vegfr expression. (F) Proposed working model. Ischemic retinal injury induces FGF2 expression in rod photoreceptors as part of an injury-response transcriptional program that promotes photoreceptor survival. In contrast, limited FGFR expression in retinal endothelial cells may constrain endothelial responsiveness to endogenous FGF2, thereby functionally uncoupling neuronal stress adaptation from pathological neovascularization.

These observations position FGF2 within a functionally associated photoreceptor stress-response network. Although our data do not establish direct transcriptional regulation of *Edn2* or *Bcl3* by FGF2, the concordant single-cell associations and bidirectional perturbation responses indicate that FGF2 is embedded within, and functionally contributes to, a broader adaptive program associated with photoreceptor survival.

### The retinal receptor landscape provides a potential cellular context for the lack of an angiogenic phenotype following FGF2 manipulation

FGF2 has well-established angiogenic activity in several experimental systems ^[31]^. The absence of a measurable angiogenic phenotype following both depletion and overexpression of photoreceptor-derived FGF2 therefore raised the question of whether the receptor context of retinal endothelial cells may contribute to this functional separation.

To explore this possibility, we analyzed the expression of FGF receptors (FGFRs) and VEGF receptors (VEGFRs) across major retinal cell populations using single-cell transcriptomic data. Retinal endothelial cells exhibited relatively limited expression of *Fgfr1-Fgfr4,* whereas canonical VEGF receptors, including *Flt1* and *Kdr,* were more prominently expressed in the endothelial compartment (Fig. 4E). In contrast, Fgfr transcripts were more evident in several neuronal populations, including photoreceptors.

This receptor distribution provides a cellular context that is consistent with the functional phenotypes observed in vivo. Although receptor transcript abundance alone cannot establish receptor activation or downstream signaling competence, the relatively limited endothelial Fgfr expression, together with the absence of vascular phenotypes following bidirectional manipulation of FGF2, suggests that endogenous photoreceptor-derived FGF2 may have limited capacity to directly engage retinal endothelial angiogenic signaling under the conditions of OIR.

Thus, the cellular distribution of ligand-producing cells and receptor expression provides a potential explanation for the functional separation between FGF2-associated neuronal stress adaptation and pathological retinal angiogenesis.

## Discussion

The present study revises our understanding of endogenous FGF2 biology during ischemic retinal injury. Rather than serving as a dominant endogenous driver of pathological neovascularization, photoreceptor-derived FGF2 functions as a stress-associated component of a neuronal adaptation program that supports photoreceptor survival under ischemic conditions. These findings address fundamental questions regarding the cellular origin, physiological function, and biological context of retinal FGF2, and provide a revised conceptual framework for understanding neurovascular responses during ischemic retinal injury.

These findings help reconcile previously conflicting observations regarding the endogenous role of FGF2 in ischemic retinopathy. Early studies showed that constitutive deletion of Fgf2 had little effect on retinal neovascularization, whereas photoreceptor-targeted FGF2 overexpression was insufficient to induce spontaneous angiogenesis ^[16]^. Conversely, subsequent studies proposed that FGF2 released from activated microglia may indirectly influence vascular remodeling through inflammatory mechanisms^[18]^. These observations may reflect the distinct cellular sources and biological contexts in which endogenous FGF2 is produced. Our rod photoreceptor-specific approach therefore helps distinguish the physiological contribution of photoreceptor-derived FGF2 from effects attributable to other cellular sources or developmental adaptation.

A key conceptual implication of our findings is that endogenous FGF2 appears to be embedded within a broader photoreceptor stress-response program rather than induced as an isolated response to vascular pathology. Previous studies have shown that photoreceptor injury induces coordinated molecular responses involving FGF2, EDN2, and other stress-associated factors, supporting the existence of conserved molecular responses to photoreceptor injury ^[22–25]^. The coordinated induction of *Fgf2, Edn2, Bcl3,* and other injury-associated genes across both OIR and MNU-induced degeneration suggests that this response represents a shared feature across distinct forms of photoreceptor injury rather than a vascular-specific program. Consistent with this interpretation, ultrastructural analysis revealed marked disruption of photoreceptor outer segments, further supporting persistent neuronal stress during OIR.

Coordinated changes in *Edn2* and *Bcl3* following bidirectional manipulation of FGF2 further support a functional link between FGF2 and this stress-response program. Although these findings do not establish direct transcriptional regulation, they suggest that endogenous FGF2 is functionally connected to a broader photoreceptor injury-response state. Importantly, this association does not imply that FGF2 is the upstream master regulator of the entire program; rather, FGF2 may represent one component of a coordinated adaptive response engaged during photoreceptor stress.

The temporal dissociation between retinal FGF2 induction and pathological neovascularization is a key feature of this injury-response program. Whereas pathological neovascularization peaked at P17 and progressively regressed thereafter^[32]^, FGF2 expression remained elevated into the regression phase, together with increased soluble FGF2 in intraocular fluid. This temporal pattern suggests that FGF2 is more closely associated with persistent neuronal stress than with the extent of ongoing vascular remodeling. More broadly, these findings highlight that neuronal injury and vascular pathology may not resolve in parallel during ischemic retinal injury, and that molecular programs associated with neuronal adaptation can persist after pathological vascular growth has begun to regress.

Genetic manipulation provided reciprocal evidence that further supports a selective role for endogenous FGF2 in neuronal resilience rather than vascular remodeling. The increased photoreceptor apoptosis following *Fgf2* deletion, together with the absence of a vascular phenotype following either deletion or overexpression, argues against the possibility that endogenous FGF2 is simply a permissive angiogenic factor whose effects become evident only when its levels are altered. Instead, these findings suggest that the principal consequence of endogenous photoreceptor-derived FGF2 in the OIR retina is related to neuronal adaptation and survival.

A central question raised by our findings is why FGF2 exhibits potent angiogenic activity in numerous experimental systems yet does not produce a measurable pathological angiogenic phenotype in the OIR retina ^[31,33]^. Our receptor landscape analysis provides a potential cellular context for this apparent discrepancy. Single-cell transcriptomic profiling demonstrated minimal expression of *Fgfr1-Fgfr4* in retinal endothelial cells, whereas the canonical VEGF receptors *Flt1* and *Kdr* were highly enriched in the endothelial compartment.

Together, these data indicate that the receptor context of retinal endothelial cells differs substantially from that of the canonical VEGF pathway. Receptor transcript abundance alone, however, cannot establish receptor activation or downstream signaling competence. Nevertheless, the concordance between the receptor expression landscape and the lack of measurable vascular phenotypes following bidirectional FGF2 manipulation supports the possibility that receptor context may contribute to the functional separation observed in vivo.

More broadly, our findings support the concept of neurovascular compartmentalization during ischemic tissue injury ^[34]^. Retinal ischemia is often viewed as a tightly coupled pathological process in which neuronal injury and vascular remodeling are intrinsically linked. Our data instead suggest that these responses can be differentially regulated and may follow partially distinct biological trajectories during ischemic retinal injury. Whereas VEGF signaling predominantly governs endothelial proliferation and pathological neovascularization ^[35]^, photoreceptor-derived FGF2 participates in an intrinsic neuronal stress-adaptation program associated with photoreceptor survival and characterized by coordinated activation of conserved stress-response genes. This framework helps explain why substantial neuronal stress persists despite vascular regression and further suggests that therapeutic strategies designed to enhance neuronal resilience could complement, rather than compete with, current anti-VEGF therapies targeting pathological vascular growth.

Several limitations should be acknowledged. First, while our study defines the cellular origin and physiological function of endogenous FGF2 in vivo, the downstream molecular mechanisms mediating its effects on photoreceptor survival remain to be fully elucidated. Future studies will be required to further define the intracellular mechanisms underlying FGF2-mediated photoreceptor protection. Second, *Edn2* and *Bcl3* were selected as representative molecular readouts because they have been implicated in photoreceptor stress responses and survival-associated pathways across retinal injury models. Thus, our study places FGF2 within this conserved stress-response network without attempting to define the complete architecture of the transcriptional program. Third, receptor expression patterns were inferred from transcriptomic data, and future studies will be needed to determine how receptor abundance and signaling competence shape FGF2 responsiveness in specific retinal cell populations. Finally, our analyses focused primarily on the acute phase of OIR, and whether photoreceptor-derived FGF2 contributes to long-term retinal remodeling or visual function requires further investigation.

In summary, we identify rod photoreceptors as the major endogenous source of FGF2 during ischemic retinal injury and demonstrate that photoreceptor-derived FGF2 functions as a component of a neuronal stress-adaptation program that supports photoreceptor survival rather than an endogenous driver of pathological neovascularization. More broadly, our findings suggest that neuronal adaptation and pathological vascular remodeling represent interconnected yet separable responses during retinal ischemia. This work not only refines our understanding of endogenous FGF2 biology but also provides a conceptual framework for integrating neuronal adaptation and vascular remodeling within the ischemic neurovascular unit, supporting future therapeutic strategies that combine control of pathological vascular growth with approaches aimed at preserving neuronal integrity.

## Materials and Methods

### Animals

*Fgf2*^fl/fl^ mice (strain S-CKO-02405) and Rho-Cre transgenic mice (strain C001018) were purchased from Cyagen Biosciences (China). Mice were maintained under a 12 h light/12 h dark cycle with free access to food and water. Rod-specific *Fgf2* conditional knockout mice were generated by crossing *Fgf2*^fl/fl^ mice with Rho-Cre mice. Littermates carrying the floxed *Fgf2* allele without Cre expression were used as controls. Genotypes were confirmed by PCR according to the manufacturer’s protocols. All experimental procedures were approved by the Institutional Animal Care and Use Committee (IACUC) of Shenzhen TOPBIOTECH Co., Ltd. (Approval No. TOPGM-IACUC-2024-0408) and were performed in accordance with institutional guidelines for animal care and use.

### OIR model

The OIR model was established as previously described ^[36]^. Briefly, P7 mice with their nursing dams were exposed to 75% oxygen for 5 days in an oxygen-controlled chamber. At P12, mice were returned to room air to induce relative retinal hypoxia and subsequent pathological neovascularization. Retinas were collected at indicated time points for molecular, histological, and vascular analyses. Quantification of pathological neovascularization was performed at P17, corresponding to the peak phase of pathological neovascularization in the OIR model.

### AAV-mediated gene delivery

To manipulate retinal FGF2 expression in a cell-type-dependent manner, recombinant AAV vectors carrying different promoters were used. For rod photoreceptor-targeted FGF2 overexpression, AAV8 vectors expressing mouse *Fgf2* under the control of the rhodopsin promoter (AAV8-Rho-*Fgf2*) were used, with AAV8-Rho-GFP serving as the corresponding control vector^[37]^.Subretinal injection was performed at P1. Briefly, 0.3 μL of viral suspension (5 × 10^12^ viral genomes/mL) was delivered into each eye using a fine glass micropipette.

For broader retinal FGF2 overexpression, AAV2 vectors carrying mouse *Fgf2* driven by the cytomegalovirus promoter (AAV2-CMV-*Fgf2*) were used, with AAV2-CMV-GFP serving as the control vector. Intravitreal injection was performed at P5 with 0.5 μL of viral suspension per eye (1 × 10^13^ viral genomes/mL). The efficiency of FGF2 manipulation was evaluated by quantitative real-time PCR using retinal tissues collected at the indicated time points.

### MNU-induced degeneration model

To induce photoreceptor degeneration, age- and sex-matched (6- to 8-week-old male C57BL/6J) mice received a single intraperitoneal injection of freshly prepared MNU (60 mg/kg, dissolved in saline with 0.1% acetic acid). Vehicle controls received the same volume of solvent. At 0, 6, and 12 h post-injection, mice were deeply anesthetized and euthanized by cervical dislocation; retinas were rapidly dissected and snap-frozen for RNA extraction. Three biological replicates (*n*=3) were used per time point.

### Single-cell RNA sequencing analysis

Publicly available single-cell RNA sequencing datasets from normoxic and OIR mouse retinas were reanalyzed in this study. Raw count matrices were obtained from the China National Center for Bioinformation (CNCB) under BioProject accession number PRJCA069552 ^[29]^ . Data processing, including quality control, normalization, dimensionality reduction, clustering, and cell-type annotation, was performed using the Seurat package (version 5.5.0) in R (version 4.4.0) ^[38]^ . Cell clusters were annotated based on the expression of canonical retinal cell-type marker genes. Expression levels of *Fgf2*, photoreceptor injury-response genes, and receptor genes (*Fgfr1-Fgfr4*, *Flt1*, and *Kdr*) were extracted and visualized across annotated cell populations.

For analysis of photoreceptor stress-response programs, rod photoreceptors were extracted based on cell-type annotation. Expression correlations between *Fgf2* and selected stress-associated genes were assessed at the single-cell level. A photoreceptor injury-response module score was calculated based on a predefined gene set consisting of *Edn2*, *Bcl3*, *Fos*, Jun family members, *Atf3,* and *Mt1.* Module scores were compared between normoxic and OIR conditions to evaluate differences in injury-response gene expression patterns.

### In situ RNA hybridization

Spatial localization of Fgf2 transcripts was performed using the PinpoRNA 2.0 in situ RNA hybridization system (PinpoRNA 2.0 kit, Cat. No. PIF2000, PinpoEase) according to the manufacturer’s instructions. Briefly, retinal cryosections were hybridized with a probe specifically targeting mouse Fgf2 transcripts (PinpoRNA probe, Cat. No. 1417311-B1). Following signal amplification and fluorescent detection, nuclei were counterstained with DAPI. Images were acquired by ZEISS LSM 980 confocal microscopy under identical acquisition settings for all experimental groups.

For quantitative analysis, cells containing at least two clearly distinguishable *Fgf2* RNA puncta within the cellular boundaries were classified as *Fgf2-* positive. Cells exhibiting only a single punctum or nonspecific background signal were not included in positive cell quantification. The proportion of *Fgf2-* positive cells was calculated as the percentage of *Fgf2-* positive cells among total DAPI-positive nuclei within the outer nuclear layer (ONL). Comparable retinal regions were analyzed across experimental groups.

### Quantitative real-time PCR

Total RNA was extracted from retinal tissues using TRIzol reagent according to the manufacturer’s instructions. Complementary DNA was synthesized using the PrimeScript RT reagent Kit (Takara, RR036A). Quantitative real-time PCR was performed using SYBR Green-based detection chemistry (Takara, RR820A). Relative gene expression levels were calculated using the comparative threshold cycle method (2^-ΔΔCt^) and normalized to the endogenous reference gene Gapdh. Primer sequences used in this study are listed in Supplementary Table 1.

### ELISA

FGF2 protein levels in intraocular fluid were quantified using a commercial mouse FGF2 ELISA kit (Elabscience, E-EL-M0170) according to the manufacturer’s protocol. Eyes were collected from 6-8 mice per sample, and intraocular fluid was collected by puncturing the cornea and sclera with a fine needle. Samples were pooled to obtain sufficient volume for ELISA measurement. Approximately 35-45 μL of pooled intraocular fluid was obtained from each sample. Samples were centrifuged at 10,000 × g for 5 min at 4°C to remove cellular debris, and supernatants were subjected to ELISA analysis. The assay was performed according to the manufacturer’s instructions, including standard curve generation, sample incubation, washing, substrate reaction, and absorbance measurement at 450 nm. Each pooled sample from 6-8 mice was considered one biological replicate, and experiments were independently repeated at least three times.

### Retinal flat-mount staining

Retinal flat-mount preparation and vascular analysis were performed as previously described^[32]^. Briefly, eyes were enucleated and fixed in 4% paraformaldehyde. Retinas were dissected, permeabilized, blocked, and stained with fluorescein-conjugated isolectin B4 (IB4) to visualize retinal vasculature. Flat-mounted retinas were imaged using ZEISS LSM 980 confocal microscopy under identical acquisition conditions. Pathological neovascularization was quantified at proper time points by measuring preretinal neovascular tuft formation according to established OIR quantification methods. The avascular area was quantified as the percentage of retinal area lacking IB4-positive vascular structures relative to the total retinal area.

### TUNEL assay

Photoreceptor apoptosis was assessed using a TUNEL apoptosis detection kit (Servicebio, G1504-100T) according to the manufacturer’s instructions. Briefly, retinal cryosections were incubated with TUNEL reaction reagents and counterstained with DAPI. Images were captured under identical exposure settings across all experimental groups. For quantification of TUNEL-positive photoreceptors in the ONL, hot spots were defined as the two most densely labeled microscopic fields per section, selected after surveying the entire ONL. For each mouse, at least three non-adjacent sections were analyzed, and the average number of TUNEL-positive nuclei per hot-spot field was calculated as one biological replicate. All quantifications were performed in a blinded manner.

### Transmission electron microscopy

For ultrastructural analysis, retinal tissues were collected and processed for TEM. Briefly, retinal samples were fixed in electron microscopy-grade fixative, dehydrated through a graded ethanol series, embedded in epoxy resin, and sectioned into ultrathin sections. Sections were subsequently stained with uranyl acetate and lead citrate before imaging. Ultrastructural images were acquired using a transmission electron microscope (HITACHI HT7800, 80 kV). Photoreceptor outer segment morphology was evaluated from TEM images obtained from the indicated experimental groups. Outer segment length was measured from the connecting cilium to the distal end of the outer segment in individual photoreceptors. Outer segment fragmentation was quantified by counting fragmented or structurally discontinuous outer segments within predefined retinal regions and expressed as the percentage of affected outer segments. Ultrastructural damage was evaluated using a predefined morphological scoring system based on the presence and severity of structural abnormalities, including outer segment fragmentation, disorganization, and other prominent ultrastructural defects. Images were analyzed using predefined criteria and, where applicable, in a blinded manner.

## Statistical analysis

All quantitative data are presented as mean ± SEM unless otherwise indicated. Statistical analyses were performed using GraphPad Prism (version 10.4.0) and R-based computational tools for single-cell RNA sequencing analyses.

For comparisons between two independent groups, two-tailed unpaired Student’s t-test was used. For comparisons among multiple groups, one-way analysis of variance (ANOVA) followed by Tukey’s multiple-comparison test was applied for datasets involving all pairwise comparisons. When comparisons were performed against a single reference group, one-way ANOVA followed by Dunnett’s multiple-comparison test was used.

For time-course experiments involving comparisons between normoxic and OIR groups across multiple time points, statistical significance at each time point was assessed using two-tailed unpaired Student’s *t*-test followed by Benjamini-Hochberg false discovery rate (FDR) correction for multiple comparisons.

For single-cell RNA sequencing analyses, gene expression correlations were assessed using Pearson correlation analysis. Cells with detectable expression of the analyzed genes were included for correlation analysis, and double-negative cells were excluded. Photoreceptor injury-response module scores were calculated based on predefined gene sets, and comparisons between experimental groups were performed using appropriate statistical tests as indicated in the corresponding figure legends.

Biological replicates (*n*) represent individual animals unless otherwise indicated. For histological and imaging analyses, measurements from multiple sections or fields within each animal were averaged and the mean value from each animal was used as one biological replicate. Statistical details, including sample sizes, statistical tests, and significance thresholds, are provided in the corresponding figure legends. A *P* value < 0.05 was considered statistically significant.

## Data availability

The single-cell RNA-sequencing datasets analyzed in this study are publicly available from the China National Center for Bioinformation (CNCB) under BioProject accession number PRJCA069552. The data supporting the findings of this study are available from the corresponding author upon reasonable request.

## Author Contributions

Zhifei Wu and Lulu Peng contributed equally to this work. Zhifei Wu, Peng Wang, and Wei Du conceived and designed the study. Zhifei Wu, Lulu Peng, Jiechun Wu, Huirong Xu, Yaling Liu, Dingqiao Wang, and Lan Wang performed the experiments and analyzed the data. Xingyu Huang performed the data analysis. Guoming Zhang provided critical guidance and manuscript suggestions. Zhifei Wu, Peng Wang, and Wei Du wrote and revised the manuscript. All authors read and approved the final manuscript.

## Acknowledgements

This work was supported by the Shenzhen Basic Research General Program (JCYJ20250604142708011), the Shenzhen Science and Technology Program (ZDSYS20220606100801004), and the Futian District Health System Research Project (FTWS2025011, FTWS2026009, FTWS2026011).

---

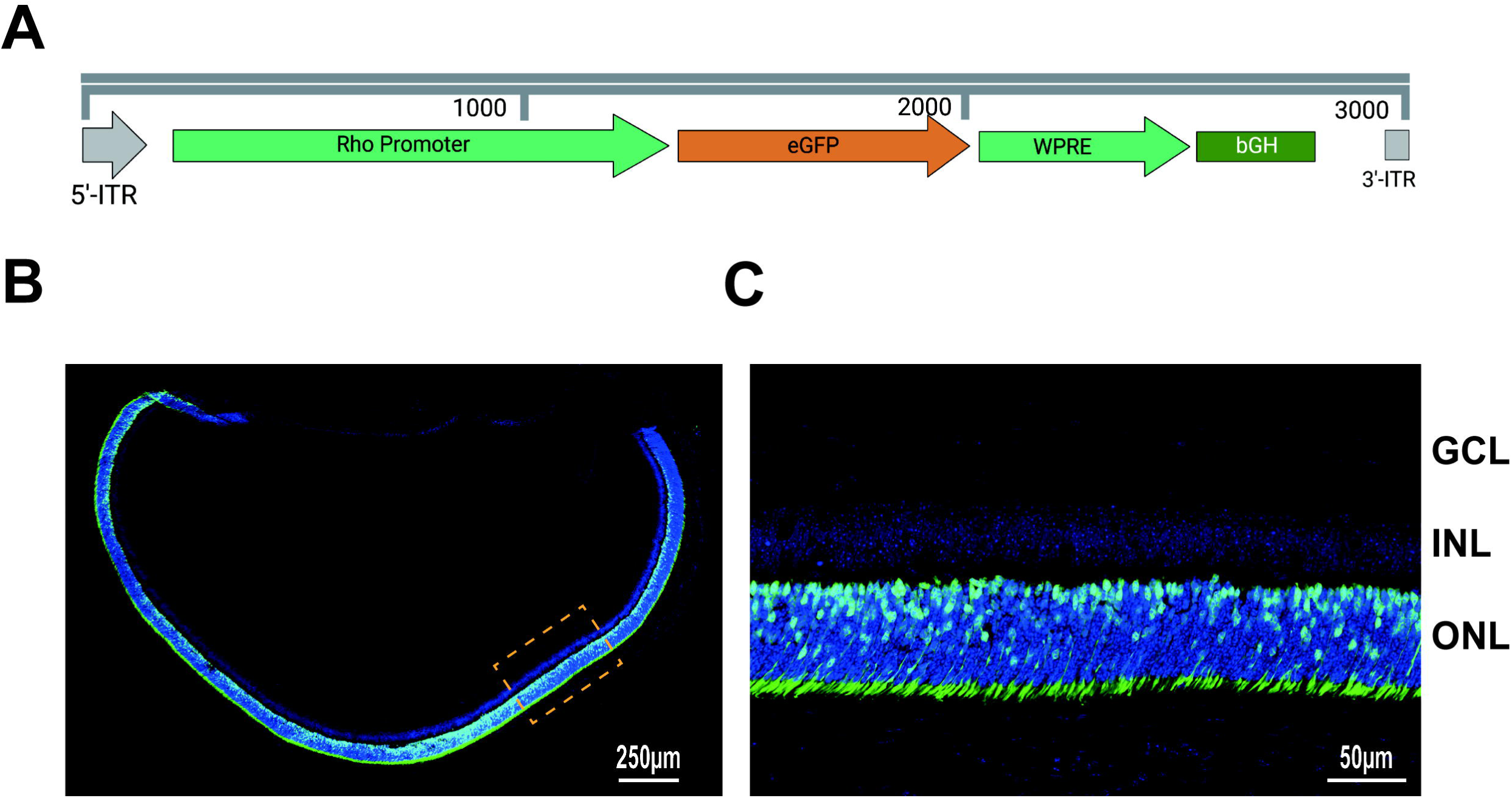

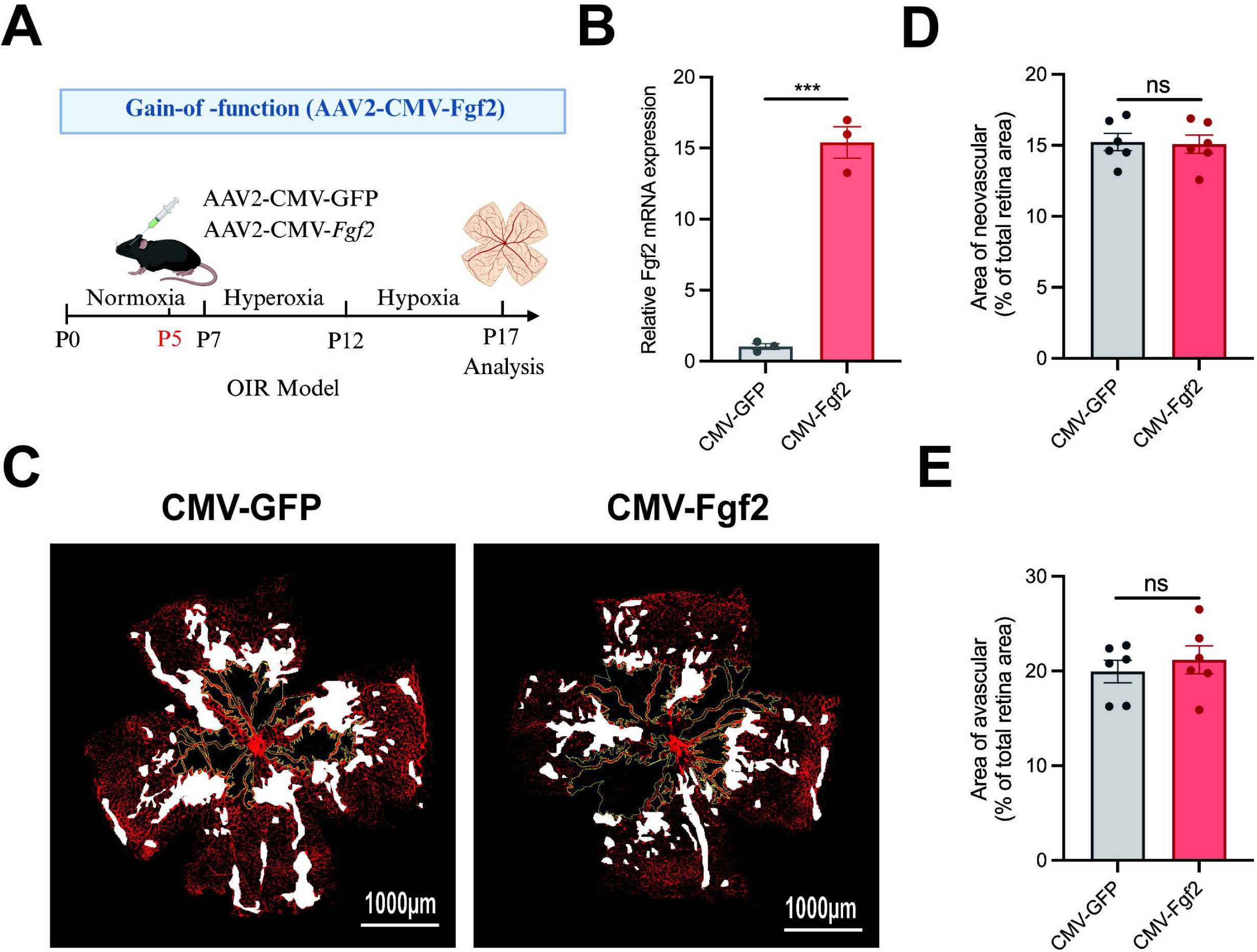

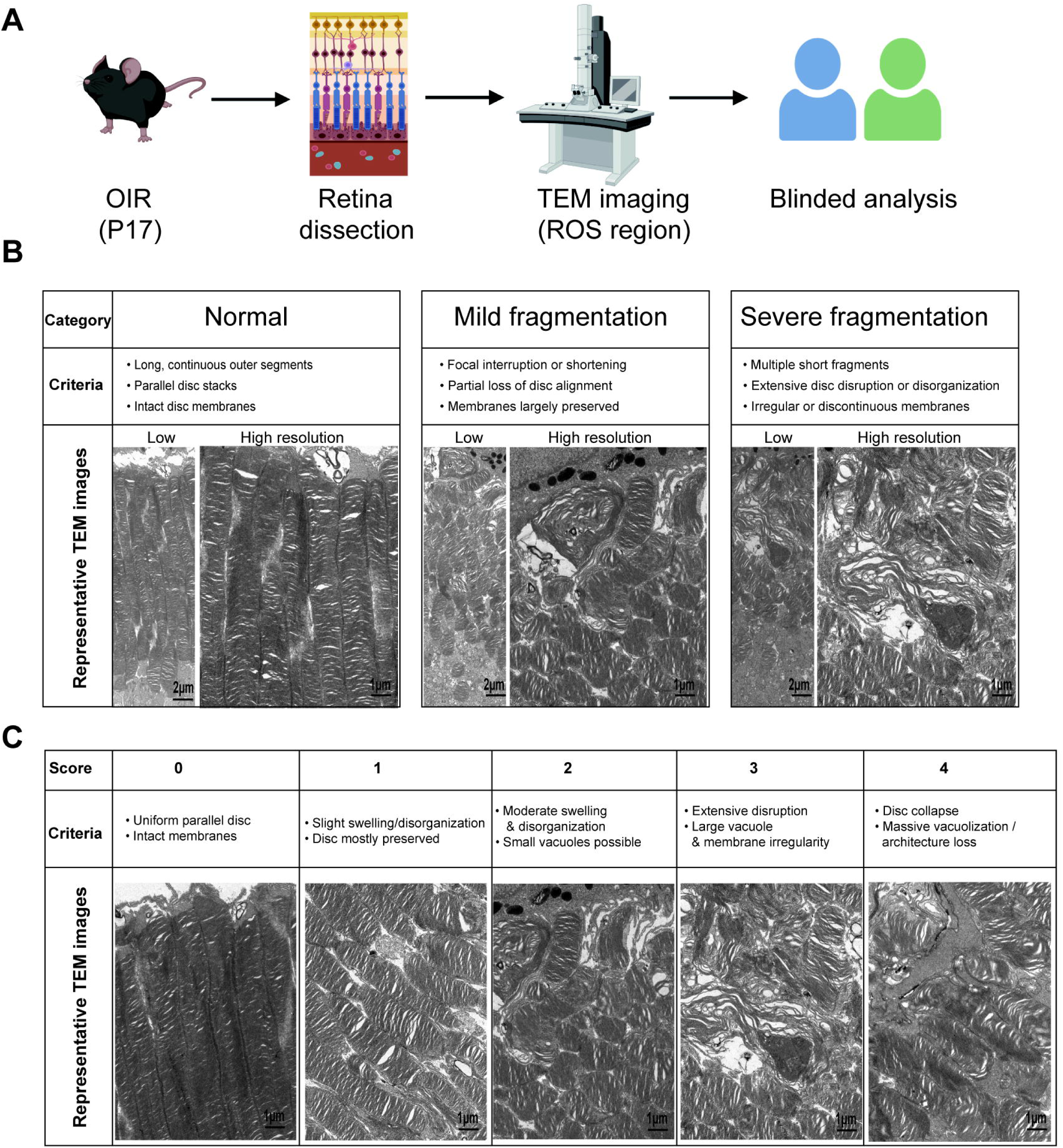

