## supplementary text for "Photoreceptor-derived FGF2 mediates a protective stress response without driving pathological retinal neovascularization in ischemic retinopathy"

**Supplementary Information**

### Table S1. Primers used for RT-PCR

| Gene | Primer sequences (5'-3') |
|---|---|
| *GAPDH*-Forward | AGGTCGGTGTGAACGGATTTG |
| *GAPDH* -Reverse | TGTAGACCATGTAGTTGAGGTCA |
| *Fgf2*-Forward | GAAACACTCTTCTGTAACACACTT |
| *Fgf2*-Reverse | GTCAAACTACAACTCCAAGCAG |
| *Edn2*-Forward | TTCTGCCATCGAAGACACTG |
| *Edn2*-Reverse | ATGGCCTTTCTTGTCACCTC |
| *Bcl3*-Forward | AGCAGTCGTCTCAGCTCCAATG |
| *Bcl3*-Reverse | AGGCAGGTGTAGATGTTGTGGG |

**Supplementary Figure Legends**

**Supplementary Figure 1. Validation of photoreceptor transduction by AAV8-Rho-GFP**

(A) Schematic representation of the AAV8-Rho-GFP construct, in which the mouse rhodopsin promoter drives GFP expression in photoreceptors.

(B) Representative whole retinal section stained with DAPI showing GFP fluorescence two weeks after subretinal injection of AAV8-Rho-GFP. GFP expression indicates successful transduction of photoreceptor cells. Scale bar, 250 μm.

(C) Higher-magnification image of the boxed region in (B), showing GFP-positive photoreceptors in the outer nuclear layer. Scale bar, 50 μm.

**Supplementary Figure 2. CMV-driven FGF2 overexpression does not alter pathological retinal angiogenesis**

(A) Schematic representation of the AAV constructs used for retinal FGF2 overexpression (AAV-CMV-*Fgf2*) and control (AAV-CMV-GFP).

(B) Quantification of retinal *Fgf2* mRNA expression by qRT-PCR following AAV-CMV-GFP or AAV-CMV-*Ffg2* injection. Data are presented as mean ± SEM; $n = 3$ mice per group. Statistical significance was determined using an unpaired two-tailed Student's *t*-test. ***$P < 0.001$.

(C) Representative IB4-stained retinal flat mounts from OIR mice injected with AAV-CMV-GFP or AAV-CMV-*Ffg2*. White-filled areas indicate pathological neovascular tufts, and yellow outlines indicate avascular areas. Scale bar, 1000 μm.

(D, E) Quantification of neovascular (D) and avascular (E) areas in OIR retinas following AAV-CMV-GFP or AAV-CMV-*Fgf2* injection. Data are presented as mean ± SEM; $n = 6$ mice per group. Statistical significance was determined using an unpaired two-tailed Student's *t*-test. ns, not significant.

**Supplementary Fig. S3. Morphological criteria for photoreceptor outer segment fragmentation and ultrastructural damage scoring**

(A) Schematic illustration of the TEM-based assessment of photoreceptor OS morphology and ultrastructural damage.

(B) Representative low- and high-magnification TEM images illustrating the classification of photoreceptor OS fragmentation. Normal OS were characterized by long, continuous outer segments with parallel disc stacks and intact disc membranes. Mild fragmentation was defined by focal interruption or shortening of the outer segments, partial loss of disc alignment, and largely preserved disc membranes. Severe fragmentation was characterized by multiple short OS fragments, extensive disc disruption or disorganization, and irregular or discontinuous membranes. Scale bars, 2 μm (low-magnification images) and 1 μm (high-magnification images).

(C) Representative TEM images illustrating the 0-4 ultrastructural damage score applied to photoreceptor OS. Score 0 was defined by uniformly parallel discs with intact membranes; score 1, slight swelling or disorganization with largely preserved discs; score 2, moderate swelling or disorganization with possible small vacuoles; score 3, extensive structural disruption with large vacuoles and marked membrane irregularity; and score 4, disc collapse with massive vacuolization and substantial loss of recognizable OS architecture. Scale bars, 1 μm.
